# Robust enhancer-gene regulation identified by single-cell transcriptomes and epigenomes

**DOI:** 10.1101/2021.10.25.465795

**Authors:** Fangming Xie, Ethan J. Armand, Zizhen Yao, Hanqing Liu, Anna Bartlett, M. Margarita Behrens, Yang Eric Li, Jacinta D. Lucero, Chongyuan Luo, Joseph R. Nery, Antonio Pinto-Duarte, Olivier Poirion, Sebastian Preissl, Angeline C. Rivkin, Bosiljka Tasic, Hongkui Zeng, Bing Ren, Joseph R. Ecker, Eran A. Mukamel

## Abstract

Integrating single-cell transcriptomes and epigenomes across diverse cell types can link genes with the *cis*-regulatory elements (CREs) that control expression. Gene co-expression across cell types confounds simple correlation-based analysis and results in high false prediction rates. We developed a procedure that controls for co-expression between genes and integrates multiple molecular modalities, and used it to identify >10,000 gene-CRE pairs that contribute to gene expression programs in different cell types in the mouse brain.

## Main text

Single-cell epigenome sequencing techniques, including snATAC-seq and snmC-seq, can identify cell-type-specific candidate *cis*-regulatory elements (cCREs), such as enhancers^1,2^. To validate putative enhancers and elucidate their function, it is important to identify the genes they directly regulate^3^. This can be accomplished by simultaneously perturbing enhancer activity and measuring gene expression in the same cells^4,5^. However, perturbation experiments are complex and to date have been used to screen pre-selected enhancers in cell types that could be cultured *in vitro*^*4,5*^. By contrast, single-cell transcriptomes and epigenomes from complex tissues, such as the brain, contain distinct genome-wide profiles from several hundred cell types^6,7^. Correlating enhancer epigenetic profiles with transcription across cell types can identify potential cell-type-specific enhancer-gene links^1,2,8^. However, genes with related functions often have correlated expression patterns, leading to incidental associations that could confound co-expression analyses with false-positives that do not reflect genuine enhancer-target gene interactions^1,2,8–10^.

To separate spurious from genuine associations, *trans* enhancer-gene correlations can be used as a negative control^11–15^. However, a principled analysis and validation of the most appropriate null model has not been performed. Moreover, different epigenetic assays, such as snATAC-seq and snmC-seq, measure distinct aspects of enhancer activity. It is unclear how the differences between these data modalities affect the sensitivity and specificity for detecting enhancer-gene correlations. Furthermore, correlation results may be strongly influenced by clustering analysis of single cell data, which in turn depends on multiple unconstrained parameters and algorithmic choices.

To address these gaps, we identify high-confidence, robust enhancer-gene links using a non-parametric permutation-based procedure to control for gene co-expression (Fig. 1a, Supplementary Fig. 1a). We first integrate single-cell transcriptomes (scRNA-seq) and epigenomes (open chromatin, snATAC-seq, and DNA methylation, snmC-Seq) to generate multi-modality profiles using a dataset with over 200,000 single cells from the mouse primary motor cortex^6^. We correlate the epigenetic state of putative enhancers with expression of nearby genes, and compare the observed correlation with two null distributions. A conventional shuffling procedure that randomly permutes cell labels effectively controls for noise present in single-cell sequencing measurements^1,2,10^. However, as we discuss below, this null distribution is confounded by gene co-expression and leads to spurious enhancer-gene associations. This challenge can be addressed statistically using generalized least squares regression^16^ (GLS), which transforms data matrices to decorrelate observations. We used a more general non-parametric approach, shuffling genomic regions to create an appropriate null distribution^11–15^. Moreover, we leveraged three complementary data modalities to cross-validate enhancer-gene links with independent data. Finally, we validated the predicted links with multimodal 3D chromatin conformation (snm3C-seq) data^17^.

**Fig. 1.**
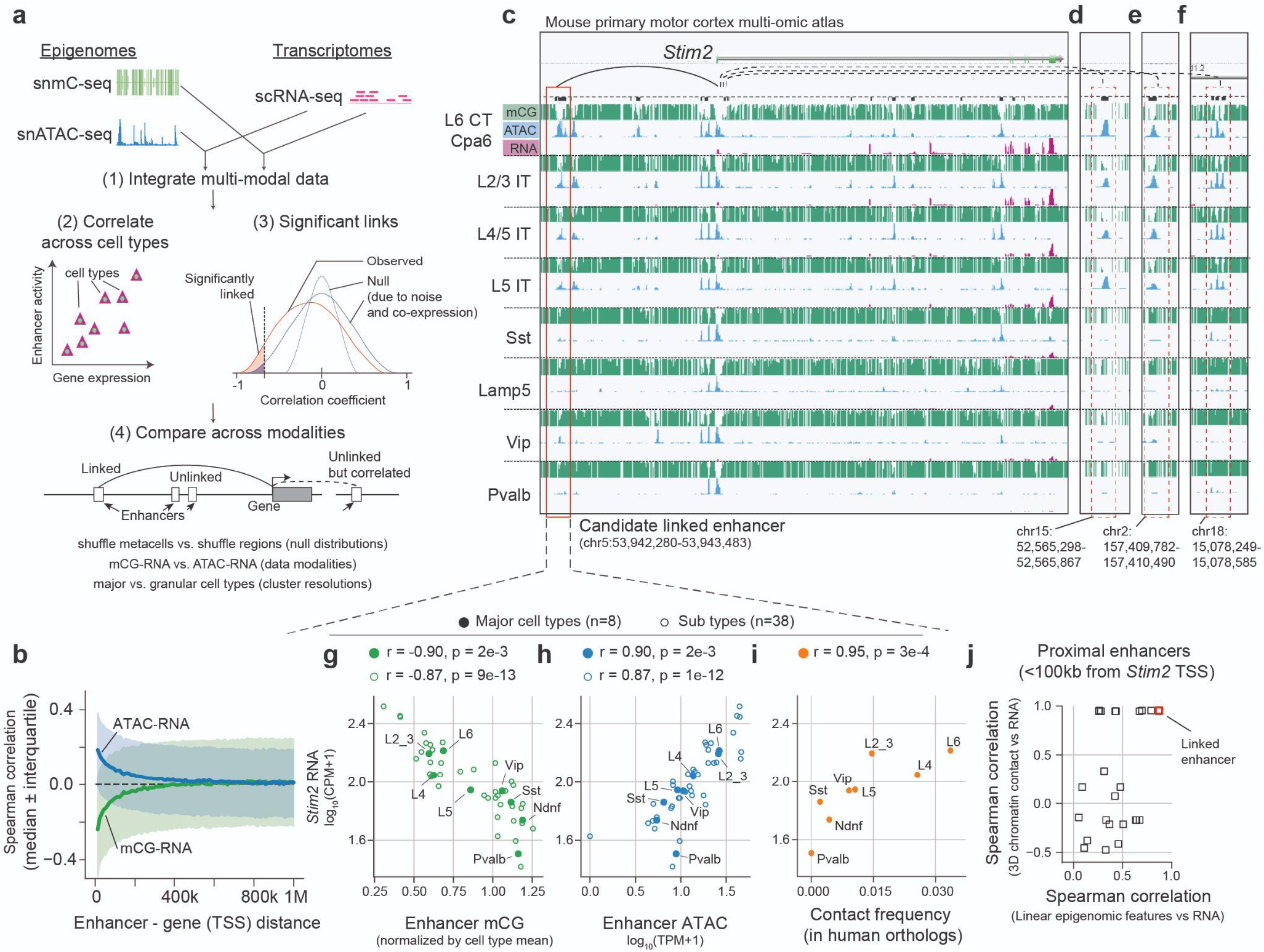
Identifying enhancer-gene links through integrated analysis of single-cell transcriptomes and epigenomes. **a**. Our proposed method links enhancers with target genes by (1) integrating single-cell transcriptomes (scRNA-seq) and epigenomes (snmC-seq and snATAC-seq), (2) correlating enhancer activity with gene expression across metacells, (3) identifying significant links compared with a shuffled null distribution, and (4) evaluating predicted links across null models, data modalities, and metacell resolutions. **b**. Strength of enhancer-gene association as a function of genomic distance. The wide interquartile range (shading) indicates high variability in enhancer-gene associations. **c-f**. Correlation of the gene *Stim2* with nearby (**c**) and distal (**d-f**) enhancer regions. **g-i**. Scatter plots of *Stim2* expression versus enhancer mCG (**g**), ATAC-seq signal (**h**), and enhancer-TSS chromatin contact frequency in human orthologs (**i**). **j**. Enhancer-gene association from linear-genome features (mCG, ATAC) versus 3D-genome features (chromatin contact frequency) for *Stim2* proximal enhancers. The x-axis shows the minimum absolute correlation value between mCG-RNA and ATAC-RNA. Enhancer mCG level is normalized by the global mean mCG level of each cell type; RNA is log_10_(CPM+1) normalized; ATAC is log_10_(TPM+1) normalized.

To illustrate the risk of false associations due to gene co-expression, we analyzed a large set of single-cell transcriptome and epigenome data from the mouse primary motor cortex^6^. Putative enhancers (see Methods; Table S1, Supplementary Fig. 1b) within ∼100 kb of a gene promoter were enriched in associations with gene expression, including positive correlations for chromatin accessibility and negative correlations for enhancer DNA methylation (mCG) (Fig. 1b, Supplementary Fig. 1c,d). However, these associations were highly variable: We observed many weak correlations for proximal enhancers (<100 kb), and relatively strong correlations for some distal enhancers (>500kb) (Fig. 1b, interquartile range ∼0.4). The broad distribution of correlation strength makes it difficult to reliably link specific enhancers with their target genes.

A representative example is the gene *Stim2*, encoding a calcium sensor that helps maintain basal Ca^2+^ levels in pyramidal neurons^18^. In cortical neurons, we identified 33 enhancers within 100 kb of the *Stim2* promoter. *Stim2* expression correlates with low mCG (r = –0.87, p=9e-13, n=38 cell types) and high chromatin accessibility (r=0.87, p=1e-12) at a nearby enhancer (Fig. 1c,g,h). By contrast, 15 other nearby enhancers have weaker, though still significant (FDR<0.05), correlation with *Stim2* expression (|r|=0.46∼0.85). Moreover, *Stim2* expression also correlated significantly with 25,027 other enhancers located throughout the genome (FDR<0.05; both mCG-RNA and ATAC-RNA), most of which (n = 23,526) were on different chromosomes (Fig. 1d-f). Such numerous correlations with *trans*-enhancers likely reflect gene co-expression, rather than direct causal links with the *Stim2* gene. For example, these *trans*-enhancers might directly regulate nearby genes whose expression patterns across cell types are similar to *Stim2* (Supplementary Fig. 1e-h,j,k).

Next, we used three-dimensional genome conformation data to test whether putative enhancer-gene links correspond to bona fide physical interactions^19^. We analyzed the 3D chromatin contact frequency of the predicted enhancer-gene pair (Fig. 1c) across homologous human brain cell types, using multi-omic snm3C-seq data^17^. Chromatin contact frequency for this enhancer was strongly correlated with *Stim2* expression (r=0.95, p=3e-4; Fig. 1i; Supplementary Fig. 1i). By contrast, other proximal enhancers were less correlated (Fig. 1j).

In addition to the challenge of widespread spurious correlations, the case of *Stim2* also illustrates the challenges associated with defining cell types^20^. For example, the same set of cells can be grouped into either 8 major types or 38 fine-grained sub-types, leading to different correlation values (Fig. 1g,h; Supplementary Fig. 1j,k).

To address these issues, we developed a procedure that controls the risk of false positives from gene co-expression, and compares predicted links across data modalities and cell type resolutions (Fig. 2a, Supplementary Fig. 2). We first integrate single-cell transcriptomes (RNA) and epigenomes (DNA methylation or chromatin accessibility) using correlated gene-level features across data modalities (SingleCellFusion)^6,21,22^. This allows us to build a neighbor graph connecting cells within and across data modalities (see Methods). Next, we define metacells^23^, which aggregate the transcriptomic and epigenomic profiles from groups of similar cells. Each metacell has a complete bi-modal (transcriptomic and epigenomic) profile, which then allows us to correlate enhancer epigenetic features with gene expression. These metacells represent cells with an adjustable resolution, capturing both discrete and continuous patterns of variation.

**Fig. 2.**
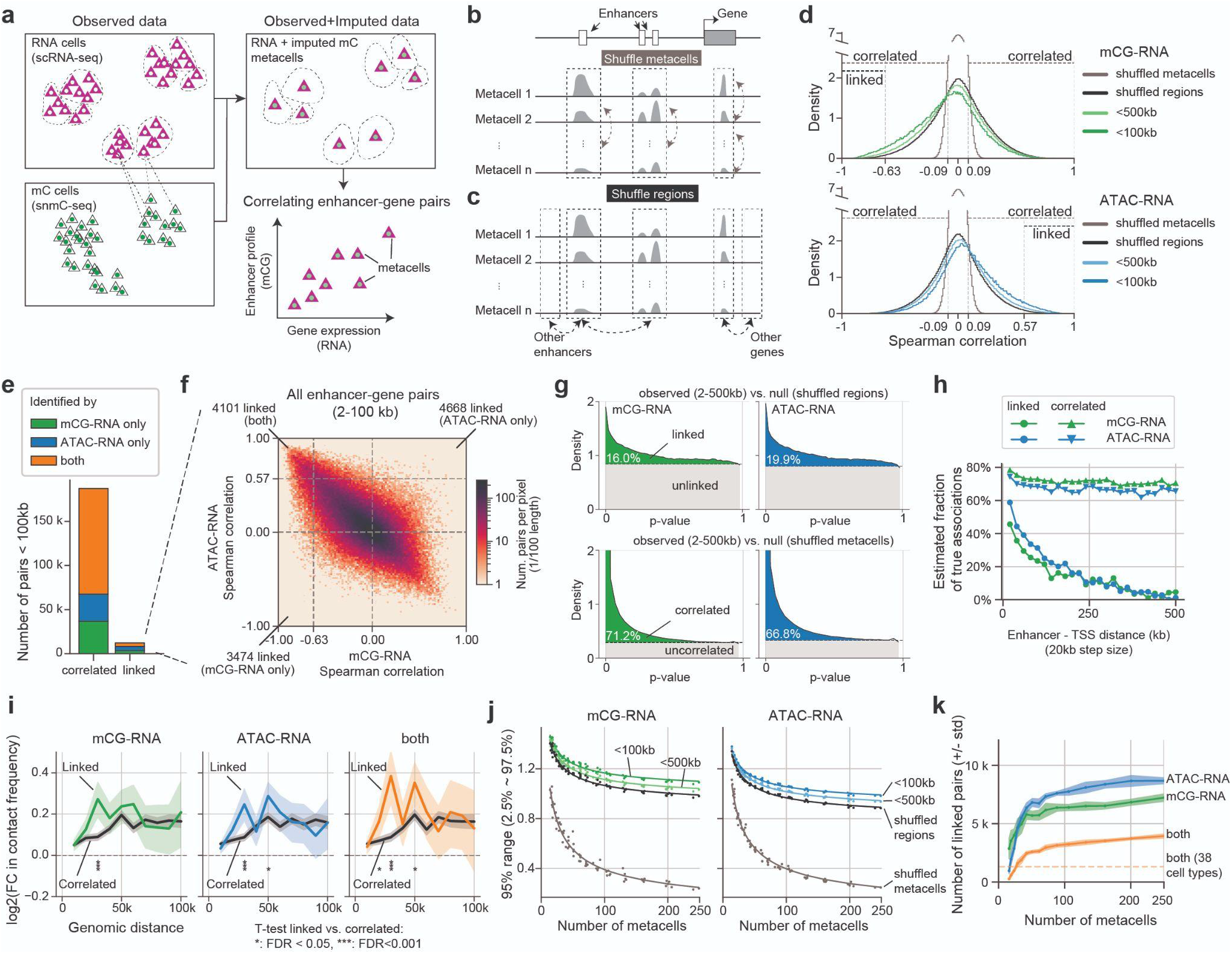
Stringent statistical criteria capture enhancer-gene links with consistent signatures across data modalities and cell type resolutions. **a**. Method for linking enhancers to target genes using metacells with bi-modality profiles. **b-c**. Null distributions derived from shuffling metacells (**b**) or shuffling regions (**c**). **d**. Distribution of enhancer-gene correlations. Bars indicate regions of statistical significance (FDR=0.2 for pairs <100kb). Two null models induce two different types of significance: linked (black bar; shuffle regions) and correlated (gray bar; shuffle metacells). **e**. The number of significantly linked or correlated pairs using mCG-RNA, ATAC-RNA, or both. **f**. Joint distribution of mCG-RNA correlation versus ATAC-RNA correlation for enhancer-gene pairs (2-100 kb). **g**. P-value histograms of enhancer-gene pairs (2-500 kb), using shuffled regions (top panels) or shuffled metacells (bottom panels). The estimated fraction of true positives is shown^24^. **h**. Estimated fraction of true associations vs. enhancer-TSS distance. **i**. Enrichment of chromatin contact frequency of linked and correlated enhancer-gene pairs compared with random genomic region pairs (mean ±95% confidence interval). Tracks are aggregated across all contacts from 8 neuronal cell types. **j**. The spread (95% range) of correlation coefficients as a function of the number of metacells. Dots represent observed data; lines represent inverse square root fit 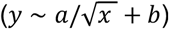. **k**. Number of linked pairs as a function of the number of metacells (FDR=0.2; mean ± standard deviation across 5 bootstrap samples with 80% of cells.)

We reasoned that genuine enhancer-gene interactions should correspond to stronger correlations than the background induced by co-expression. Correlations mediated by co-expression are inherently limited in their strength by the magnitude of gene-gene correlations, whereas direct enhancer-gene interactions can produce stronger associations. Importantly, this assumption applies to the strongest enhancer-gene interactions; weak interactions that don’t exceed the background of gene co-expression cannot be detected by correlation-based methods.

To test whether the observed correlations exceed what is expected due to noise and gene co-expression, we compared the observed correlation coefficients with two null distributions: shuffling metacells^1,2,10^ and shuffling regions^11–13^ (Fig. 2b-d). Shuffling metacells decouples epigenetic and transcriptomic signatures across metacells, removing both enhancer-gene correlation and gene co-expression (Fig. 2b). The significance arising from this distribution is inflated by gene co-expression, potentially leading to false positives in which an enhancer-gene pair may be correlated due to shared upstream regulation rather than direct interaction. Shuffling regions retains the gene co-expression structure imposed by the hierarchical organization of cell types, but it correlates each gene’s expression with distant, randomly selected enhancers (Fig. 2c)^11–13^.

As expected, the distribution obtained by shuffling regions was wider than that derived from shuffling metacells (Fig. 2d), reflecting incidental correlations due to gene co-expression. Enhancer-gene pairs within 500kb of the TSS are significantly enriched in both positive and negative correlations when compared with shuffling metacells. However, when compared with shuffling regions, enrichment is only present in positive correlation for ATAC-RNA, and in negative correlation for mC-RNA. Thus, shuffling regions is a more stringent null distribution for calling significant enhancer-gene links, as it effectively controls for spurious enhancer-gene correlations due to gene co-expression.

We call an enhancer-gene pair significantly “correlated” if it passes an FDR-adjusted threshold based on shuffling metacells, whereas we reserve the term significantly “linked” for pairs that pass the criteria set by shuffling regions. We used a relatively lenient FDR threshold of 0.2 to reduce the risk of false negatives from our stringent null distribution. Linked pairs (n=12,243 within 100kb, FDR<0.2) are a subset of correlated pairs (187,343 within 100kb, FDR<0.2) (Fig. 2e,f), but they have a stronger association that rises above the background from gene co-expression. Lowering the FDR threshold to 0.1 or 0.05 reduced the number of linked pairs to 3,142 and 489, respectively.

Notably, we found that removing sample covariance using GLS abolished the difference between shuffling regions and shuffling cells (Supplementary Fig. 3a-b). This manipulation thus removes the distinction between correlated pairs and linked pairs (Supplementary Fig. 3c). In addition, the shuffling-regions null distribution was robust with respect to differences in enhancer GC content and an enhancer’s distance to its nearest gene (Supplementary Fig. 4a-d).

We compared our results with two alternative strategies for estimating enhancer-gene interactions using single-cell epigenomes. Using open chromatin data, CICERO^8^ identified 1,869 significant enhancer-gene associations located within 100kb. These significantly overlap with a subset of the correlated pairs we identified, and to a lesser degree with linked pairs (Supplementary Fig. 5a,b). Notably, the mean CICERO co-accessibility scores are 4.8-5.9 fold higher (p<2e-8) for linked pairs than for correlated pairs (Supplementary Fig. 5c). A second strategy, the activity-by-contact (ABC) model^5^, identified enhancer-gene links using both chromatin accessibility and chromatin conformation data. This model identified enhancer-gene links for each cell type independently, without considering correlated variability in expression across cells. The ABC model identified 150,228 associations within 100kb, which significantly overlap with our correlated and linked pairs (Supplementary Fig. 5d,e). In addition, the ABC scores are 1.09-1.22 fold higher (p<1e-8) for linked pairs than for correlated pairs (Supplementary Fig. 5f). These results show that linked pairs have stronger associations than correlated pairs, and are more likely to capture genuine enhancer-gene associations.

A potential pitfall of our stringent enhancer-gene linking procedure is a higher risk of false-negatives, i.e. failure to detect true interactions. We next empirically compared correlated versus linked pairs from several biological and statistical perspectives, to test whether the correlations filtered out by our method are likely false positives arising from gene co-expression.

First, we observed that correlated pairs include many enhancer-gene links with a non-canonical direction of association (Fig. 2d; Supplementary Fig. 6a). For example, we found about a third (47,137/150,285) of these pairs had a negative correlation of gene expression with chromatin accessibility, and a similar proportion (53,687/156,932) had a positive correlation with mCG. Non-canonical associations were also reported in recent large-scale studies of brain cell epigenomes^1,2^. These correlations could suggest novel biological mechanisms such as methylcytosine-preferring transcription factors^25^. However, they may also include false-positive associations due to gene co-expression. Indeed, none of the non-canonical associations passed our threshold for linked pairs (Fig. 2d). This is consistent with the canonical understanding of enhancer activity associating with low DNA methylation and open chromatin.

Second, as enhancer-gene interactions are mostly concentrated within ∼100-500 kb around gene promoters^4,5^, we compared the distance dependence of linked and correlated pairs. Using a p-value histogram method^24^, we estimated 16.0-19.9% of enhancers that are 2-500kb away from a promoter are linked (Fig. 2g, Supplementary Fig. 6b). A much larger fraction (66.8-71.2%) were correlated. Notably, the proportion of correlated pairs remains high even for distal pairs (e.g. >60% for pairs >1 Mb or on other chromosomes), whereas <5% of these pairs are linked (Fig. 2h, Supplementary Fig. 6c). These correlated pairs contradict the biological understanding that most enhancers activate genes in *cis*; the linked pairs are more coherent with this canonical framework.

Third, we validated our predicted links with independent chromatin conformation data from the human brain^17^. We reasoned that linked enhancer-gene pairs which are conserved across species should have higher chromatin contact frequency compared with random regions. Indeed, we found enrichment of contact frequency for both linked (mean fold change (FC) = 1.15, p=2e-4) and correlated pairs (mean FC = 1.10, p=1e-5). Moreover, linked pairs located 10-30 kb apart have higher levels of contact enrichment than correlated pairs (FDR<0.05; Fig. 2i, Supplementary Fig. 6d,e).

A key parameter for our analysis is the cell type granularity, as determined by the number of metacells. The sparse genomic coverage of single-cell sequencing and the limited number of profiled cells create a tradeoff between the number of metacells and the quality of each metacell--i.e. between fine-grained resolution and signal/noise ratio. As the number of metacells (*N*) increases, the width of the null distribution for the shuffled metacells approaches zero as 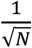, which is consistent with independent random signals for each metacell (see Methods; Fig. 2j; Supplementary Fig. 7a-c). By contrast, the range of the null distribution for shuffled regions does not vanish for large *N*, but instead asymptotes at a non-zero value that reflects gene co-expression (Supplementary Fig. 7c). Notably, the shuffling-regions null distribution is less sensitive to the number of metacells, and more closely reflects the behavior of the observed correlations. This suggests enhancer-gene link calling using shuffling-regions is less sensitive to the choice of cell type granularity than using shuffling-metacells. We found more linked pairs as the number of metacells increases, but with diminishing returns after *N* > 50. (Fig. 2k; Supplementary Fig. 7d).

We used our procedure to comprehensively examine regulatory interactions in neurons of the mouse primary motor cortex^6^. Linked enhancer-gene pairs formed 15 modules that capture diverse cell-type-specific signatures (Fig. 3a,b). For example, genes in module 13 are specifically expressed in pan-inhibitory neurons, with corresponding low CG methylation level and open chromatin at linked enhancers. Module 9 is most active in caudal ganglionic eminence (CGE) derived inhibitory neurons (Lamp5, Sncg, and Vip) and in superficial-layer excitatory neurons (L2/3 IT and L4/5 IT). These consistent gene- and enhancer-level signals integrated from three data modalities provide strong support for our identified enhancer-gene associations.

**Fig. 3.**
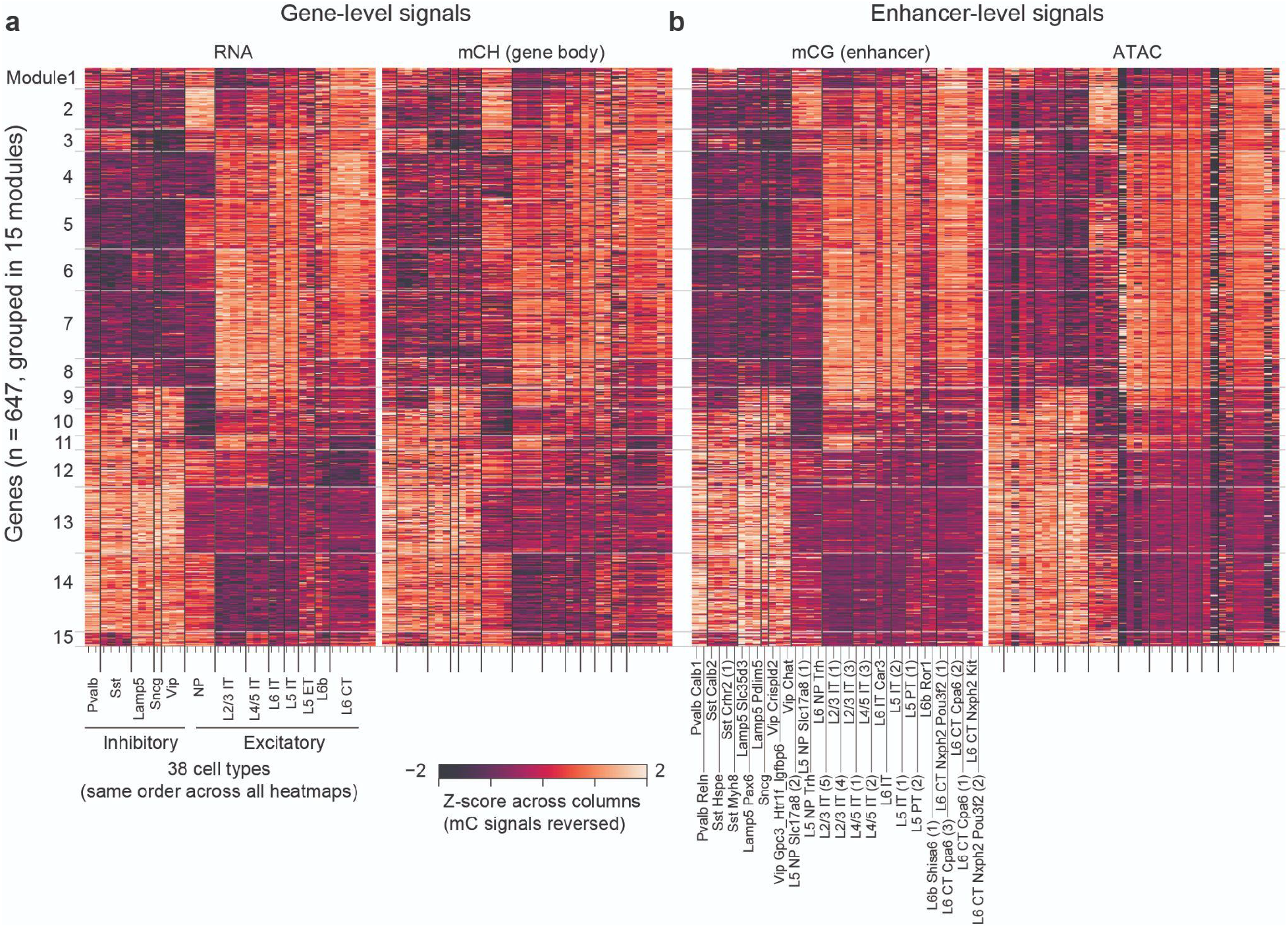
Consistent gene- and enhancer-level signatures for hundreds of enhancer-gene links. **a-b**. Gene expression (**a**), gene body DNA methylation (**b**), and enhancer mCG (**c**) and ATAC signal (**d**) across cell types. Genes are organized into 15 modules by K-means clustering. Enhancers are ordered according to the genes they are linked to (FDR < 0.2 for both mCG-RNA and ATAC-RNA across n=38 cell types). Signals from multiple enhancers linked to the same gene were averaged. The colormap for the mC modalities (gene body mCH and enhancer mCG) are reversed.

Our analyses highlight the challenge of distinguishing genuine enhancer-gene interactions from spurious correlations due to gene co-expression. We addressed this by empirically estimating the expected correlations for unlinked enhancer-gene pairs under co-expression, and comparing results across different epigenetic assays and cell type granularities. Notably, mCG-RNA and ATAC-RNA associations show striking similarities (Fig. 2d,f-k; Fig. 3), despite measuring distinct epigenetic features with opposite effects on gene expression. Predicted enhancer-gene links are robust with respect to a wide range of cell type granularities (Fig. 2k). We identified hundreds of genes and thousands of linked cCREs with highly coordinated gene- and enhancer-level activities (Fig. 3a,b).

Correlation-based analysis has notable limitations. First, this approach cannot identify constitutive enhancer-gene links that are present in all cell types. Larger datasets including more diverse tissues or cell types may partly address this limitation. Second, rigorous control for spurious correlations limits the power of detecting genuine but weak enhancer-gene interactions. Finally, true causal interactions cannot be inferred from correlational analysis alone. The links we identified (Fig. 3a,b) are strong candidates for causal enhancer-gene interactions, which must be tested by perturbative experiments^26,27^. Future experimental validation, including large-scale assays^4,5,28^, will be needed to test correlation-based predictions. By bringing together multiple data modalities to define robust enhancer-gene links, these analyses can reveal the regulatory principles of cell-type-specific gene expression.

## Supporting information

Table_S4_cell_type_correspondence

Table_S3_linked_atac

Table_S2_linked_mc

Table_S1_enhancers

## Supplementary Figures

**Supplementary Figure 1.**
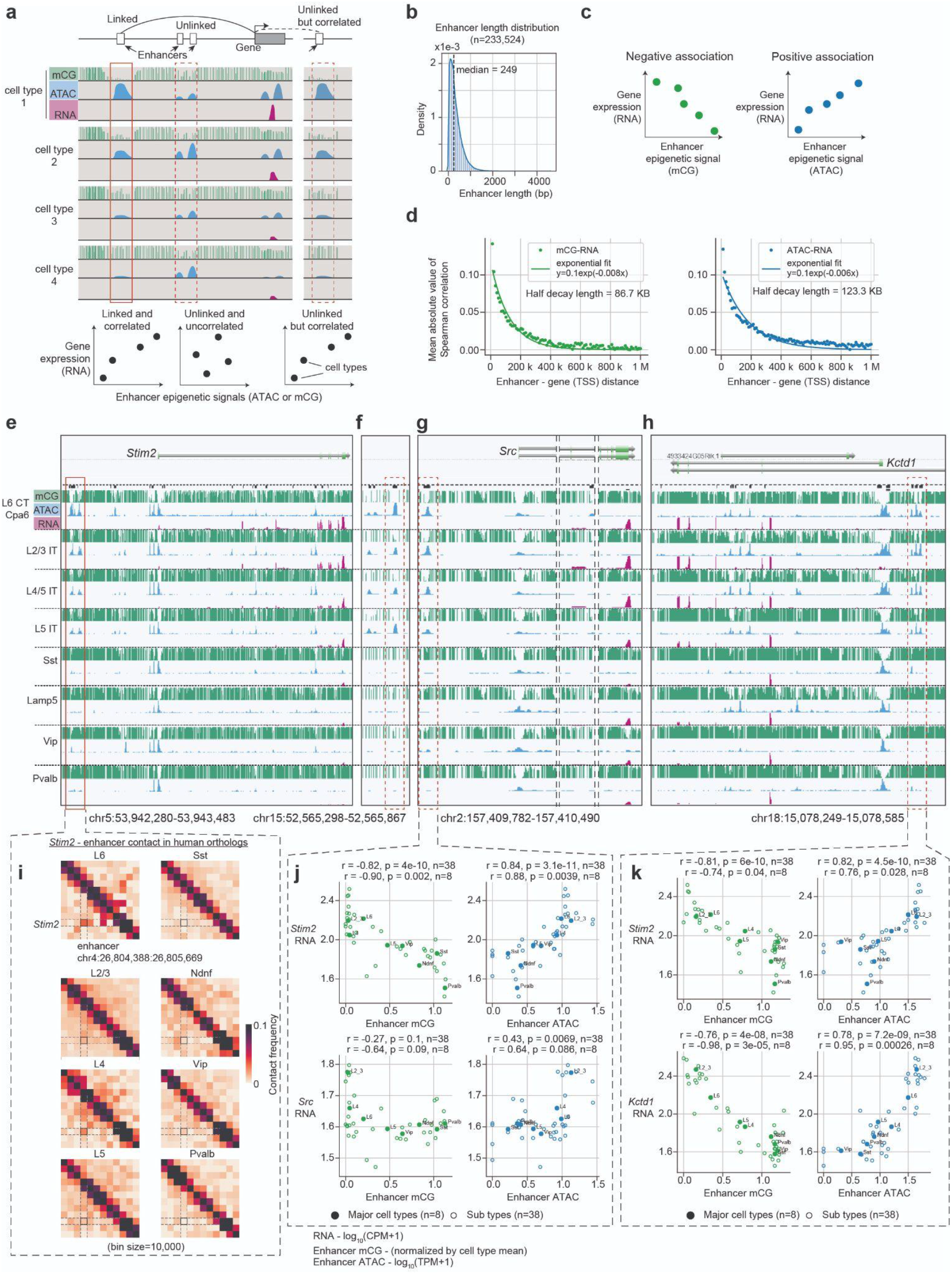
Examples of enhancer-gene links (Related to Fig. 1). **a**. Approach for linking enhancers to target gene(s) by correlating enhancer activities and gene expression across cell types. Statistically significant correlation alone may not distinguish genuine vs. spurious links. **b**. Distribution of putative enhancer length (list adapted from Ref^6^; see Methods). **c**. Illustration of two modes of enhancer-gene associations: enhancer mCG typically have positive correlation with gene expression, while enhancer ATAC-seq signals typically have negative correlation. **d**. Median Spearman correlation as a function of enhancer-TSS distance. Both mCG-RNA and ATAC-RNA decay exponentially, with a half decay length of 86.7 kb and 123.3 kb, respectively (Related to Fig. 1b). **e-h**. Genome browser views across cell types and data modalities near the gene *Stim2* (**e**), as well as other regions (**f-h**) with strongly correlated enhancer signals. Note that the highlighted enhancers in (**g-h**) are also correlated with the expression of their nearby genes (*Src* and *Kctd1*) (Related to Fig. 1c-f). **i**. Heatmaps of chromatin contact frequency in human brain cells near *Stim2* and the human ortholog of the highlighted enhancer across 8 human neuronal cell types. **j-k**. Scatter plot of *Stim2* expression (upper row) / local genes expression (lower row) versus the highlighted enhancers. Enhancer mCG level is normalized by the global mean mCG level of each cell type; RNA is log10(CPM+1) normalized; ATAC is log10(TPM+1) normalized.

**Supplementary Figure 2.**
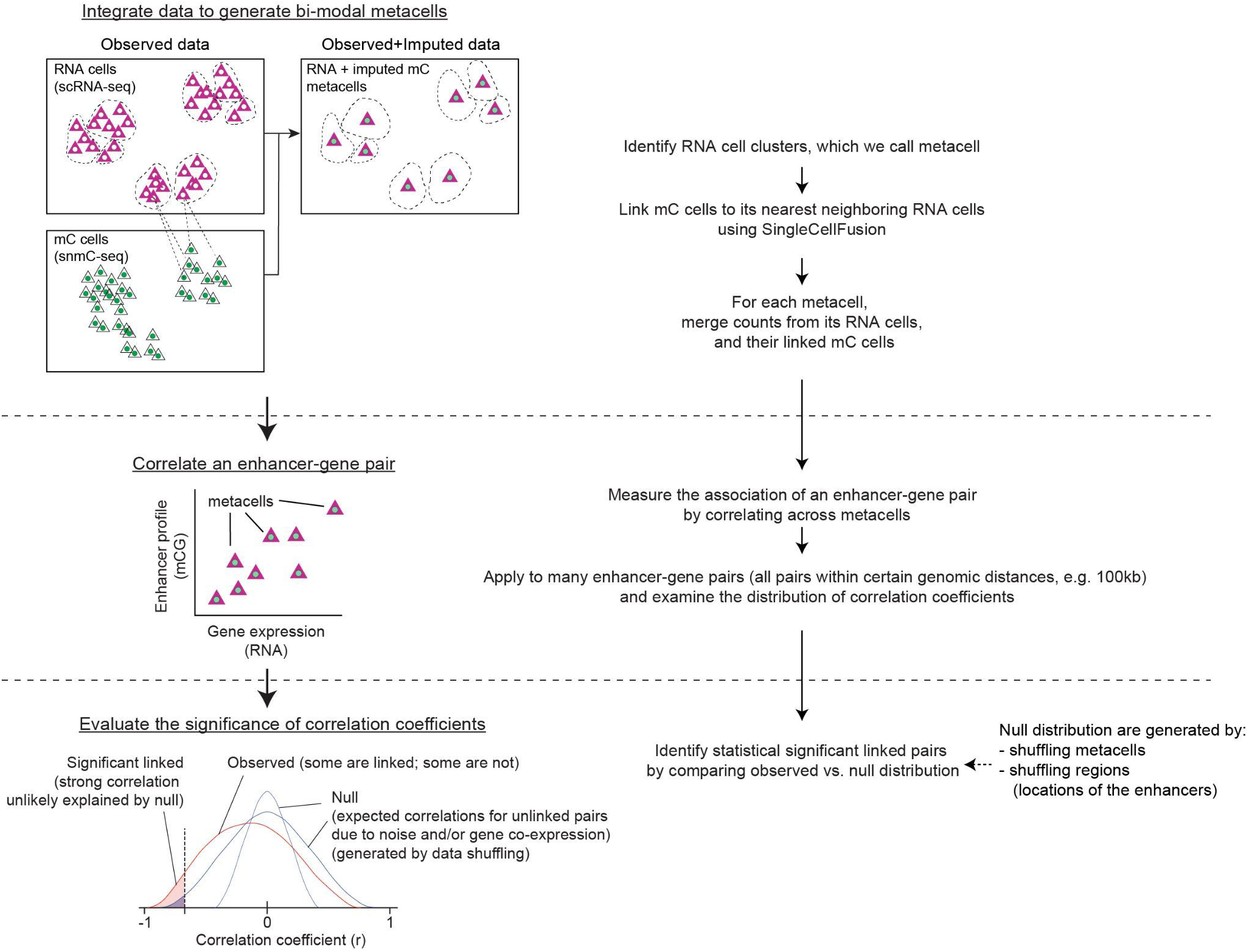
Method overview. The analysis involves three main steps. 1. Integrate transcriptomic and epigenomic data to generate metacells with bi-modal profiles. 2. Correlate enhancer-gene pairs to get correlation coefficients for individual enhancer-gene pairs. 3. Evaluate the statistical significance of correlations by comparing the observed correlations with null distributions generated by data shuffling.

**Supplementary Figure 3.**
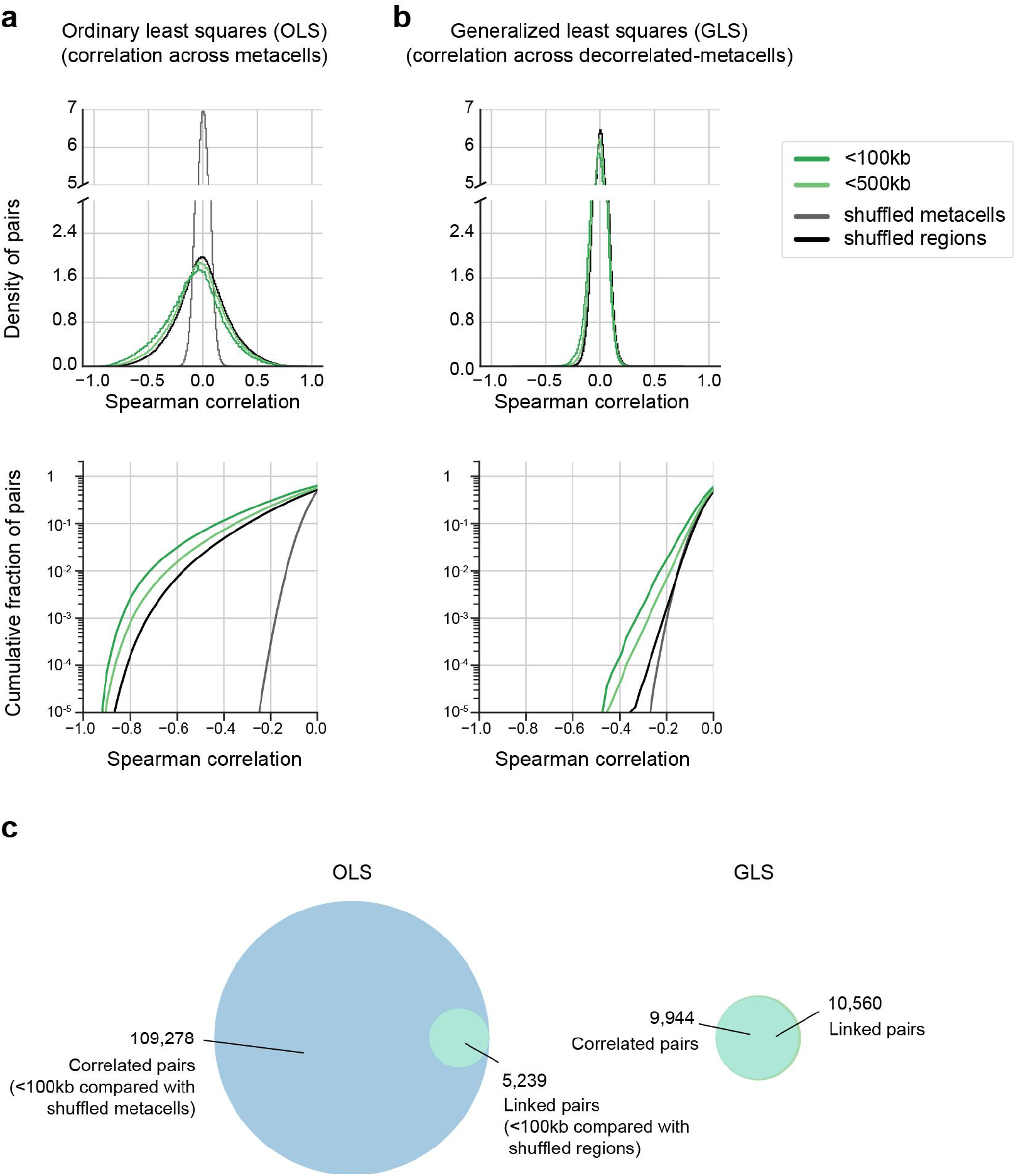
Generalized Least Squares (GLS) transformation abolishes the difference between shuffling metacells and shuffling regions (Related to Fig. 2). **a-b**. Density distribution (top) and cumulative distribution (bottom) of enhancer-gene correlations across metacells using OLS (**a**), and across decorrelated metacells using GLS (**b**). GLS transformation decouples the covariance across metacells, making the shuffling-regions distribution similar to the shuffling-metacell distribution (see Methods). **c**. Venn diagram showing the degree of overlap between correlated pairs and linked pairs, using OLS (left) and GLS (right) approaches. GLS abolishes the difference between correlated and linked pairs.

**Supplementary Figure 4.**
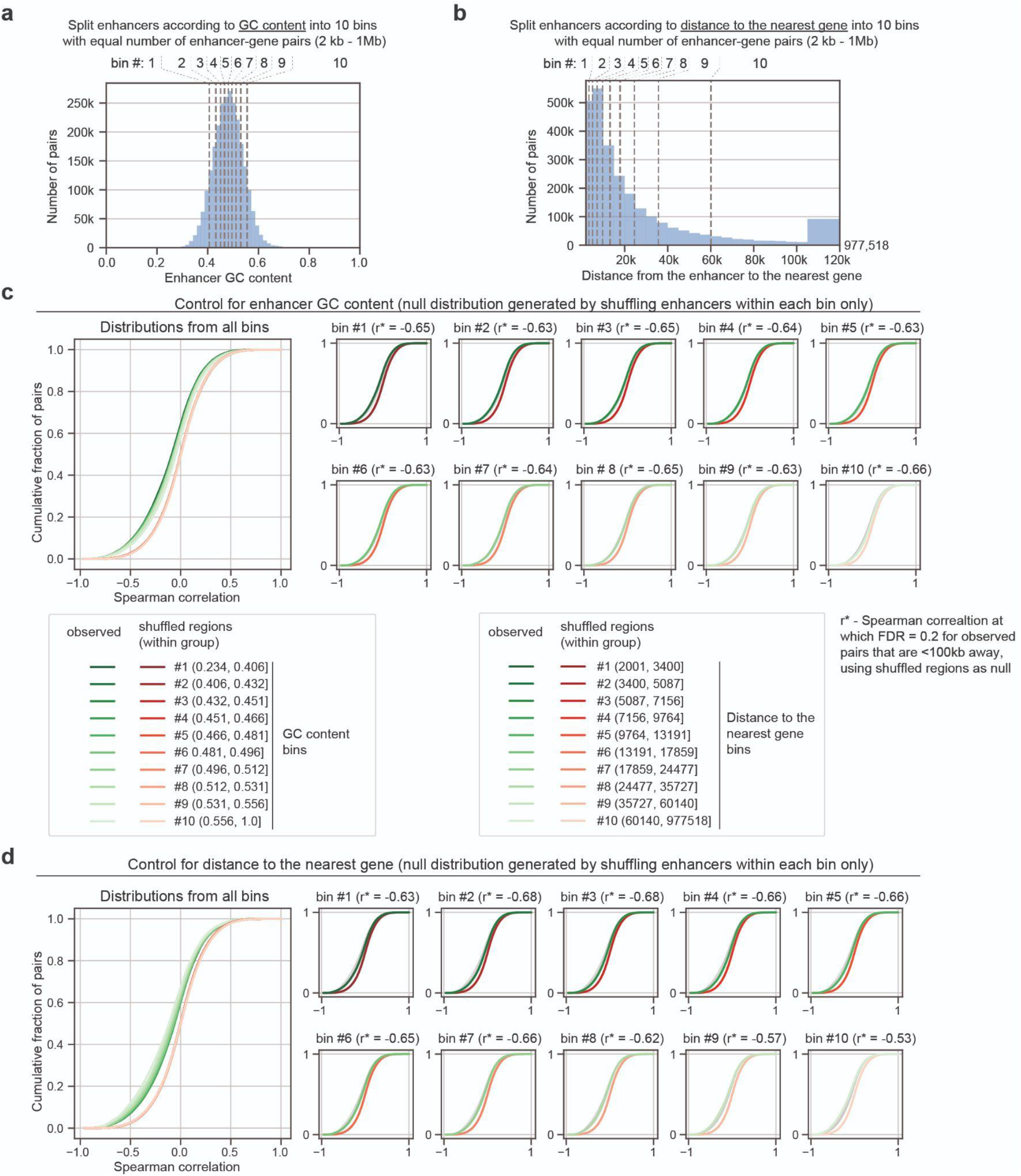
The shuffling-regions null distribution is robust with respect to enhancer GC content and distance to the nearest gene (Related to Fig. 2). **a-b**. Distribution of GC content (**a**) and distance to the nearest gene (**b**) for enhancers that are in all enhancer-gene pairs (2kb - 1Mb). In each case, they are grouped into 10 bins (deciles) with an equal number of enhancer-gene pairs. **c-d**. Cumulative distribution of enhancer-gene correlation (mCG-RNA; observed (<100kb) vs. null (shuffling regions)). The same analyses are applied to each of the 10 bins by enhancer GC content (**c**) and by distance to the nearest gene (**d**), respectively. Null distributions from different bins highly overlap.

**Supplementary Figure 5.**
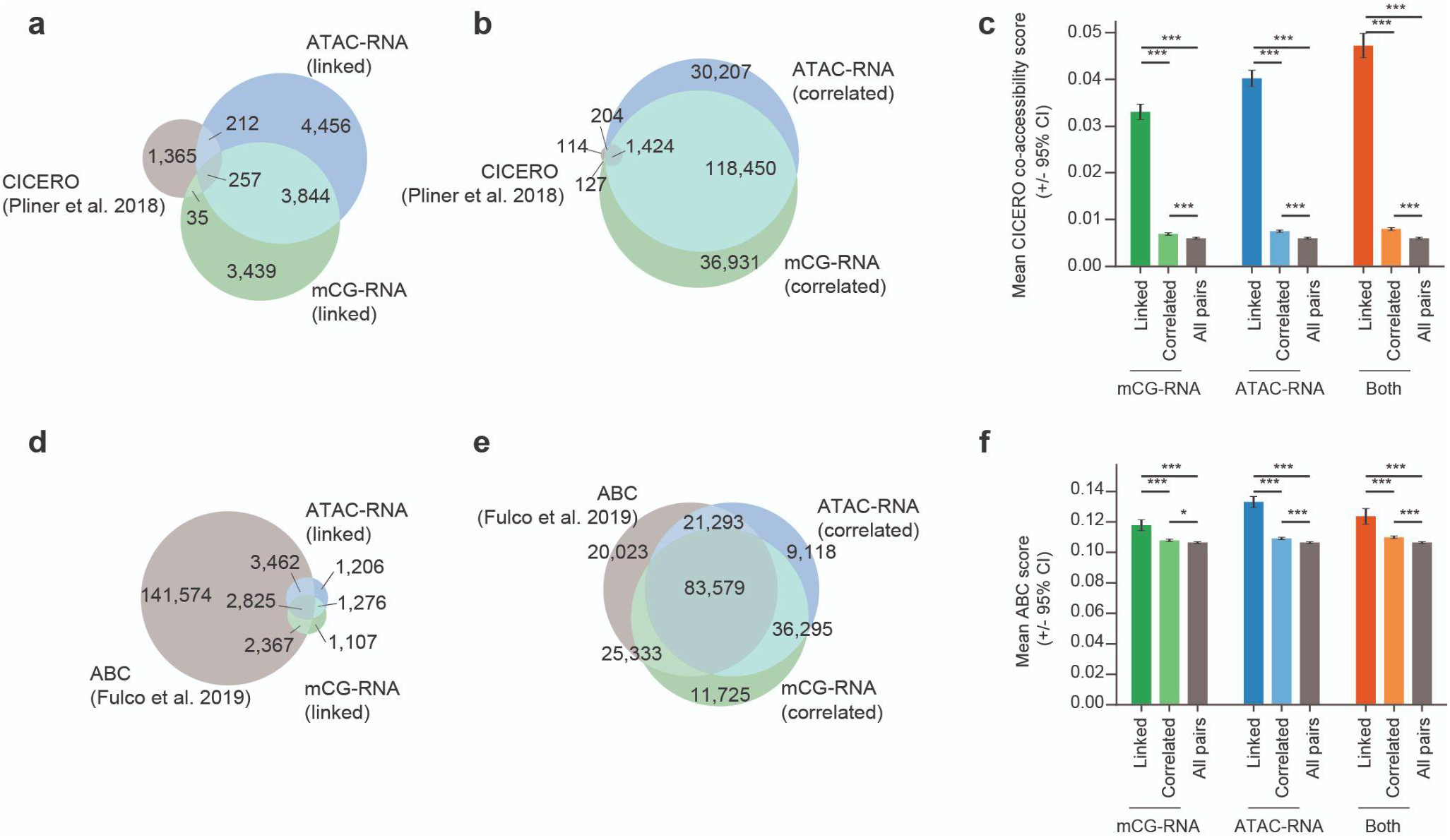
Comparison with CICERO^9^ and the activity-by-contact (ABC) model^5^ (Related to Fig. 2). **a-b**. Venn diagram comparing the enhancer-gene associations identified by applying CICERO^8^ to the mouse MOp data^6^ versus linked pairs (**a**) and correlated pairs (**b**) found in this study. **c**. Barplots comparing the mean CICERO scores across different groups of enhancer-gene pairs identified in this study. Error bars indicate 95% confidence intervals. Independent t-test are used to compare between groups (* p<0.05, *** p<0.001). **d-e**. Venn diagram comparing the enhancer-gene associations identified by applying ABC model^5^ to the mouse MOp data^6^ versus linked pairs (**d**) and correlated pairs (**e**) found in this study. **f**. Barplots comparing the mean ABC scores across different groups of enhancer-gene pairs identified by this study. ABC scores are generated for each enhancer-gene pair and cell type (n=38). We first took the maximum across cell types, followed by taking the mean of each group of enhancer-gene pairs. Error bars indicate 95% confidence intervals. Independent t-test are used to compare between groups (* p<0.05, *** p<0.001).

**Supplementary Figure 6.**
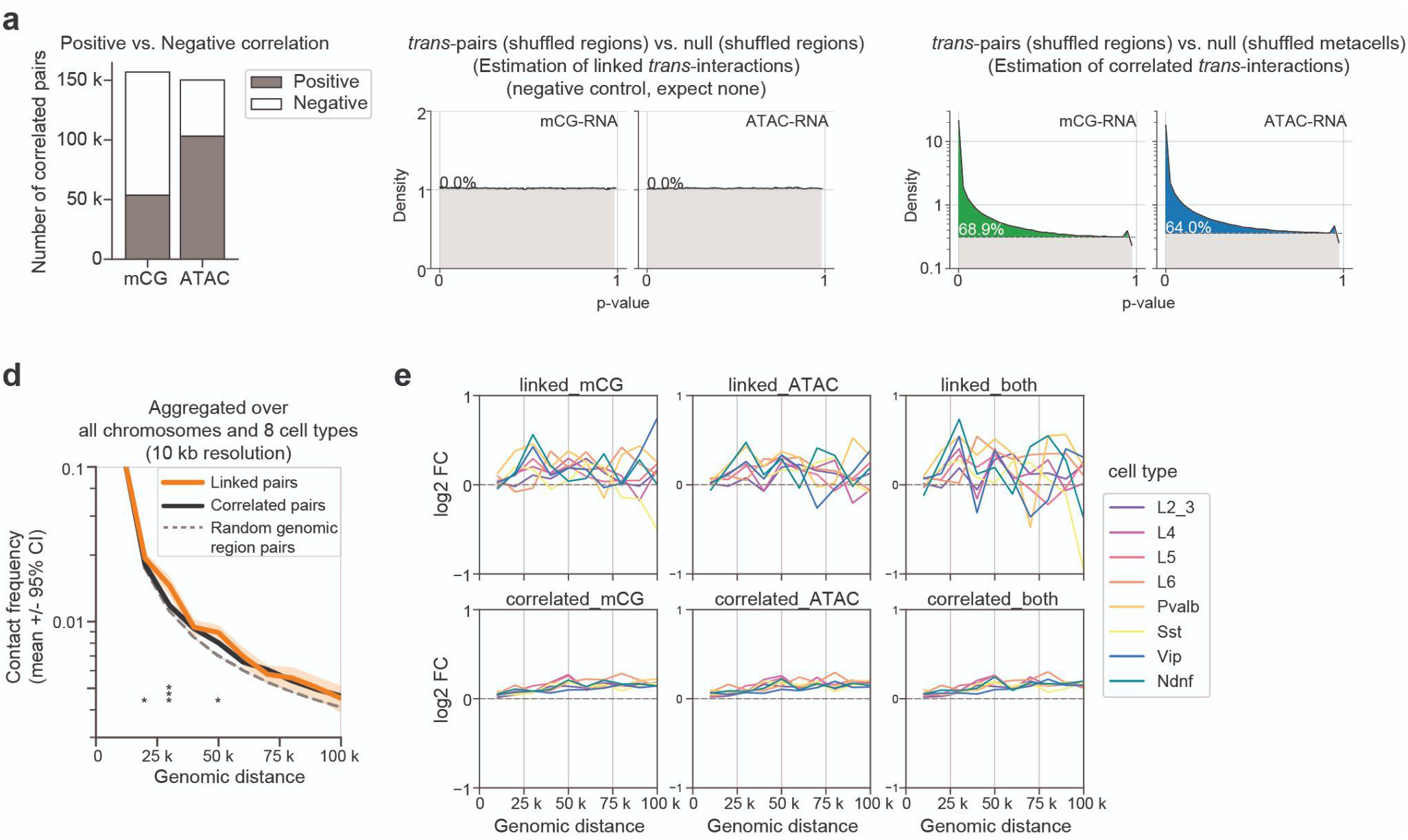
Linked vs. correlated enhancer-gene pairs have distinct characteristics (Related to Fig. 2). **a**. The number of positively or negatively correlated enhancer-gene pairs for mCG-RNA and ATAC-RNA, respectively. **b**. P-value histograms^24^ of *trans*-enhancer-gene pairs using shuffling-regions as the null distribution. The histogram closely follows a uniform distribution, indicating trans-enhancer-gene pairs are linked. This serves as a negative control for Fig. 2g. **c**. P-value histograms^24^ of *trans*-enhancer-gene pairs, using shuffling-metacells as the null distribution. The numbers mark the fraction of p-values that deviate from the uniform distribution, which estimates the fraction of correlated *trans*-enhancer-gene pairs. **d**. Chromatin contact frequencies of pairs of genomic bins as a function of genomic distance. Linked and correlated pairs (lifted over from mm10 to hg38^29^) are compared with random genomic pairs. Results are aggregated over all chromosomes (autosomes + chrX) and 8 different human neuronal cell types (L2/3, L4, L5, L6, Pvalb, Sst, Vip, Ndnf) at 10kb resolution of chromatin contact maps^17^. **e**. Enrichment of contact frequency of linked and correlated enhancer-gene pairs compared with random genomic region pairs across 8 human neuronal cell types.

**Supplementary Figure 7.**
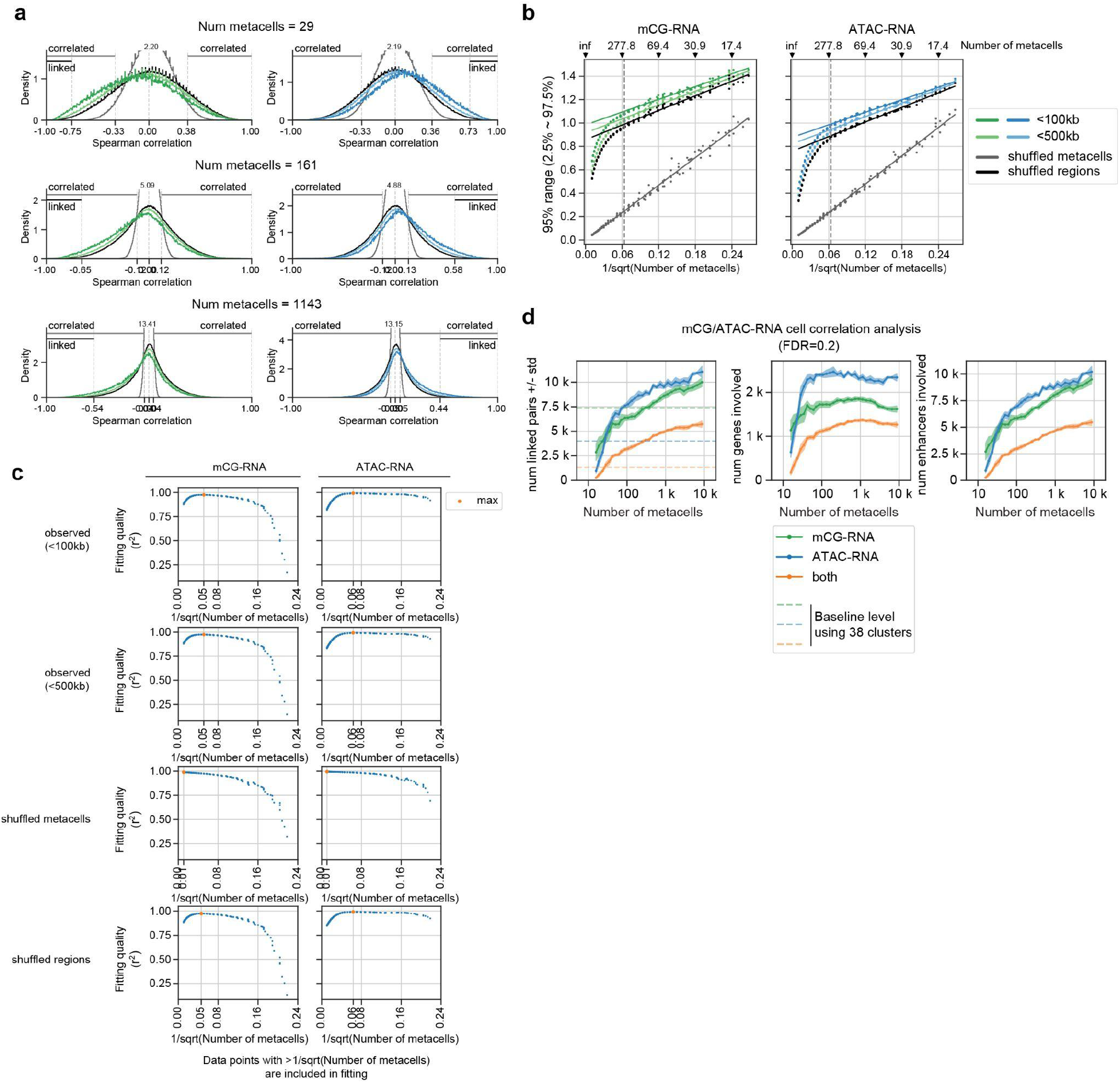
Effect of the granularity of metacells on enhancer-gene correlations (Related to Fig. 2). **a**. Distributions of correlation coefficients for different numbers of metacells. The distributions become narrower as the number of metacells increases. Here the number of metacells controls cell type granularity. **b**. Range of correlation coefficients (2.5%∼97.5% range) as a function of 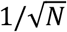, where *N* is the number of metacells. Data points are well fitted by a straight line for *N* < 277. **c**. Fitting quality, as measured by *r*^2^, as a function of fitting cutoff--range of data points in (**b**) used for fitting. The fitting quality peaks at 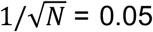, i.e., *N*= 278. **d**. The number of linked pairs (left), number of genes involved (middle), and number of enhancers involved (right) as a function of the number of metacells.

## Methods

### Datasets

We used three single-cell sequencing datasets from the mouse primary motor cortex (MOp)^6^. They are scRNA-seq (single cell; 10x genomics V3; Allen Institute for Brain Science), snmC-seq (single nucleus; DNA methylation; Ecker lab from the Salk Institute), and snATAC-seq (single nucleus; chromatin accessibility; Ren lab from UCSD). Only high-quality neuronal cells, as determined in Ref^6^ (from its <u>Supplementary Table 2</u>; column SCF/SingleCellFusion), are retained for our analysis. These datasets are publicly available and provided by a previous study (Ref^6^; https://assets.nemoarchive.org/dat-ch1nqb7). The starting point of all analyses are gene-by-cell matrices and/or enhancer-by-cell matrices depending on the data modality. For the scRNA-seq dataset, we start from the gene-by-cell count matrix. For the snATAC-seq dataset, we quantified both enhancer-by-cell and gene-by-cell count matrices. For the snmC-seq dataset, we quantified enhancer-by-cell CG DNA methylation profiles and gene-by-cell non-CG (CH) DNA methylation profiles. The DNA methylation profile for a particular region and cell can be summarized by two numbers: the number of methylated cytosines (mC) and the total number of cytosines covered (C). The DNA methylation level is the ratio of mC to C (mC/C). Please see sections below for dataset specific procedures of normalizations. The mouse gene annotation file is downloaded from gencode (vM16). The enhancer list is adapted from the putative enhancer list from Ref^6^ (see below).

### Calling putative enhancers

We constructed our putative enhancer list based on the mouse MOp neuronal cell type-specific putative enhancers from Ref^6^ (from its <u>Supplementary Table 7</u>). In that study, the enhancers are called using REPTILE^30^, an algorithm that uses the DNA methylation and ATAC-seq profiles of 13 mouse neuronal cell types, as well as mouse embryonic stem cells, as input. Starting from this list, we first selected regions with enhancer score >0.5 and merged overlapping regions using bedtools^31^. We subsequently removed regions overlapping any gene promoter regions (transcription start site +/- 2kb; all transcripts from gencode vM16), exons (vM16), and ENCODE blacklist^32^. This leaves us with 233,524 enhancers in total, with a median size of ∼250 bp (Supplementary Fig. 1b; Table S1).

### Curated cell types

For analyses related to Figure 1, we curated a list of 38 neuronal cell clusters based on the SingleCellFusion clusters (L1 and L2, with n=29 to 56 cell types respectively) in Ref^6^. We aimed to merge small clusters to increase pseudo bulk coverage at enhancers, while retaining as much cell type diversity as possible. To achieve this, we first call an enhancer *covered* in a cluster if it has at least 20 sequenced CpG sites in that cluster, where the cluster-level coverage is the sum of cell-level coverages. Next, we call an enhancer *common*, if it is covered in more than half of the L2 clusters. We call a cluster *covered*, if more than half of the common enhancers are covered in that cluster. For each L1 cluster we then evaluate 3 cases:

1. If the cluster itself is not covered, we drop it along with all its child (L2) clusters.
2. Else if less than 2 (n<2) of its child (L2) clusters are covered, we retain the L1 cluster itself, but drop all its child (L2) clusters.
3. Else if at least 2 (n≥2) of its child (L2) clusters are covered, we retain the covered L2 clusters, but drop the uncovered L2 clusters and the L1 cluster.

This procedure resulted in 38 clusters with adequate coverage. Table S4 summarized the correspondence between the 38 clusters we get from this procedure and the cell types defined in Ref^6^.

To compare with the cell types in snm3C-seq data^17^, we further merged these 38 fine-grained clusters into 8 major clusters based on the well-established neuronal cell type taxonomy^33^. Table S4 summarized the correspondence between the 38 fine grained and the 8 major cell clusters defined in this study and those defined in Ref^6,17^.

### Clustering and defining metacells

For analyses related to Figure 2, we generated cell clusterings with a range of cluster resolutions. We start by normalizing the scRNA-seq count matrix with log_10_(CPM+1), where CPM stands for counts per million mapped reads. We then calculated the top 50 principal components (PCs), and built a k-nearest neighbor graph (k = 30) connecting cells according to the Euclidean distance in the PC space. We used Leiden community detection to generate clusters^34^. Different resolution parameters (r = 1 ∼ 794) were chosen to generate clusters with different granularity (n = 13 ∼ 8850 metacells). The pseudo bulk profiles from each of the individual clusters were used as metacells.

### Feature selection and normalization

We preprocessed the data matrices separately for each data modality. The starting point is always cell-level matrices containing counts (RNA and ATAC) or methylation level (mC). To get cluster-level (metacell) matrices, we summed counts from cells in the same clusters (metacells) to create pseudo-bulk samples. For methylation data, we summed methylated counts and total counts (coverage) separately. Next, we normalized matrices as follows:

- For an RNA matrix (gene-by-cluster/metacell), we normalize the raw count matrix with log_10_(CPM+1).
- For an ATAC matrix (enhancer-by-cluster/metacell), we normalize the raw count matrix with log_10_(TPM+1), where TPM stands for transcripts per million mapped reads. Enhancers that are covered in <50% of clusters are removed.
- For a gene body mCH matrix (gene-by-cluster/metacell), we first removed low coverage genes if the gene has <50% clusters surpassing 1000 counts in the gene body (or < 80% metacells surpassing 20 counts). We then take the ratio of the number of methylated to the number of coverage to get the methylation fraction. All the steps here consider cytosines in non-CG (CH) dinucleotide context only.
- For an enhancer mCG matrix (enhancer-by-cluster/metacell), we first removed low coverage enhancers if the gene has <50% clusters surpassing 20 counts (or <80% metacells surpassing 5 counts) in the enhancer region. We then take the ratio of the number of methylated to the number of coverage to get the methylation fraction. All the steps here consider cytosines in CG dinucleotide context only.

After normalization and filtering of individual matrices, we then consider only enhancers that are shared in both ATAC and mCG matrices for downstream analyses.

### Correlating enhancer-gene pairs across cell types

We calculate the Spearman correlation coefficient between any pair of enhancer and gene that are within 1 Mbp (enhancer center to gene TSS) across curated cell types (n=38 or n=8). This was done separately for enhancer mCG vs. RNA and enhancer ATAC vs. RNA. Enhancer mCG signals are normalized by the global mean mCG levels of each cell type; enhancer ATAC signals are log_10_(TPM+1) normalized; RNA expression levels are log_10_(CPM+1) normalized.

To assess the statistical significance of the enhancer-gene correlations, we repeated the correlation analysis with 2 types of data shuffling control, as explained in the main text. To control for random noise, we shuffled cell cluster labels of the gene-by-cluster RNA matrix, followed by calculating correlation coefficients. To control for background co-expression across enhancer-gene pairs, we shuffled gene labels of the gene-by-cluster RNA matrix, followed by calculating correlation coefficients.

### Correlating enhancer-gene pairs across metacells

Given a transcriptomic dataset (scRNA-seq) and an epigenetic dataset (e.g. snmC-seq) collected from the same tissue, we first generate a constrained k-nearest neighbor network linking cells across the two modalities (SingleCellFusion; Ref^6,22^). This network allows us to impute the DNA methylation profiles (mC) for each RNA cell. We then cluster scRNA-seq cells using Leiden community detection ^34^ (see section **Clustering/Generating metacells**). We call these clusters *metacells*, to emphasize that they do not necessarily correspond to discrete cell types, but could also capture continuous changes among cell populations. These preparations allow us to construct bi-modal profiles for each metacell, by aggregating counts--either observed or imputed--from cells in the same metacells. Finally, we evaluate the correlations between enhancer-gene pairs across metacells.

To be specific, the starting point of this analysis involves 4 matrices: an enhancer-by-cell mCG (or ATAC) matrix *E*_*ec*_, a gene-by-cell RNA matrix *R*_*gc*_^*′*^, a cross-modal cell-to-cell k nearest neighbor matrix: *K*_*cc′*_, and a metacell assignment matrix of RNA cells *K*_*c*_^*′*^_*z*_. Here we use *ci c′*and *z* to denote an mC cell, an RNA cell, and a metacell, respectively. A metacell is a group of RNA cells generated by Leiden clustering. We use *g* and *e* to denote an enhancer and a gene, respectively. All matrices contain unnormalized raw counts. *K*_*cc′*_ is generated by SingleCellFusion^6,22^ with default settings and cross-modal k=30. *K*_*c′z*_ is generated by Leiden clustering on the RNA-seq dataset as mentioned in previous sections.

To get bi-modal profiles for a metacell, we aggregate counts from the cells belonging to that metacell: *R*_*gz*_ = ∑_*c′*_ *R*_*gc′*_*K*_*c′z*_, and *E*_*ez*_ = ∑_*c*_ *E*_*ec*_*K*_*cc′*_*K*_*c′z*_. The metacell profiles are then normalized as mentioned in previous sections to adjust for metacell size, library size, and gene length. Finally, normalized *R*_*gz*_ and *E*_*ez*_ allow us to correlate a specific pair of gene *g*^(*i*)^ and enhancer *e* ^(*i*)^ across metacells (*z*). We calculated Spearman correlation coefficients for all enhancer-gene pairs with distance between 2kb to 1Mb (enhancer center - TSS).

### Estimating the statistical significance of enhancer-gene links

To assess the statistical significance of a correlation coefficient *r*, we constructed two null distributions by shuffling metacells (Fig. 2b) and shuffling regions (Fig. 2c). In the first case, we shuffle metacell labels independently for transcriptomic and epigenetic data, such that the two data modalities become independent of each other. In the second case, we permute and enhancers randomly from their original genomic location to the locations of other genes and enhancers, while retaining the linked bi-modal profiles of each metacell.

Either null distribution can be used to get empirical p-values and false discovery rate (FDR). The empirical p-value of a correlation coefficient *r* is defined as the cumulative fraction of the null distribution that has more extreme (stronger) correlation coefficients than *r*. We calculated two-sided p-values when using the shuffled metacells distribution, and single-sided p-values when using the shuffled regions distribution. FDRs are then calculated using the Benjamini-Hochberg procedure^35^. We call an enhancer-gene pair significantly *linked* (*correlated)* if its empirical FDR is <0.2 using shuffling regions (metacells) as the null.

To see if the shuffled regions distribution depends on enhancer properties such as its sequence GC content and distance to the nearest gene, we also performed stratified shuffling analyses (Supplementary Figure 4). We first grouped enhancers into 10 bins (deciles) according to their GC content or distance to the nearest gene. We then shuffled enhancers within each bin and compared observed enhancer-gene correlations with shuffled ones for each bin separately.

### Enrichment of 3D chromatin contact frequencies

We validated the predicted enhancer-gene links using single-cell measurements of 3D-chromatin contact frequency in human prefrontal cortex ^17^. Raw contact matrices of 8 neuronal cell types were downloaded as mcool files ^17^. We calculated contact frequencies from raw counts using matrix balancing using Cooler^36,37^. We then focused on analyzing these contact frequency matrices at a resolution of 10kb non-overlapping genomic bins across the genome.

To compare our enhancer-gene links predicted in the mouse brain with the chromatin contact data from human brain, we lifted genes (gencode vM16 whole genes) and putative enhancers from mm10 to hg38 using LiftOver^29^ with parameters -minMatch=0.8 and - minBlocks=1.00.

To calculate enrichment, we first assigned enhancers (center) and genes (TSS) to their corresponding genomic bins (non-overlapping 10kb bins genomewide). We compared the contact frequencies of the predicted enhancer-gene pairs with random genomic region pairs with similar genomic distance. We separately tested the enrichment of contact frequencies of 6 groups of predicted enhancer-gene pairs: mCG-RNA linked, ATAC-RNA linked, pairs linked by both modalities, mCG-RNA correlated, ATAC-RNA correlated, and pairs correlated in both modalities. For each of the 8 neuronal cell types, we only include pairs that are active in the specific cell type, i.e. whose gene expression is greater than the median across all 8 cell types.

### Comparison with CICERO

We installed the R package CICERO^8^ from the Bioconductor following the instructions from the authors’ tutorial (https://cole-trapnell-lab.github.io/cicero-release/docs_m3/#constructing-cis-regulatory-networks). We ran CICERO on MOp ATAC-seq data using default parameters. The program takes as input a peak-by-cell ATAC-seq matrix, where peaks include both putative enhancers we specified and gene promoters (500 bp upstream of TSS). The program returns co-accessibility scores for peak pairs. We filtered the output down to enhancer-promoter pairs only, removing enhancer-enhancer and promoter-promoter pairs. We also focused on analyzing enhancer-gene pairs that are within 100kb apart, to compare with our correlation-based analysis. We used a threshold = 0.2 following Ref^9^ to call positive enhancer-gene pairs.

### Comparison with the ABC model

We downloaded code from the github repository of the ABC model^5^ (https://github.com/broadinstitute/ABC-Enhancer-Gene-Prediction) and followed instructions. We ran ABC for each MOp cell type (n=38) using our identified putative enhancer list (n=233,524) and pseudo-bulk ATAC-seq and RNA-seq data as input. We used genomic-distance based power law estimation to model chromatin contacts (--score_column powerlaw.Score). The software returns a score (ABC score) for each enhancer-gene pair and cell type. We excluded the expressed genes from the results, as suggested by the authors. We also focused on analyzing enhancer-gene pairs that are within 100kb. We used a threshold = 0.022 as recommended by the authors to call positive enhancer-gene pairs.

### Generalized least squares (GLS) analysis to decouple covariance across metacells

We used GLS ^16^ to test the association between gene expression and enhancer activity across cell types (metacells). We will focus on only one given enhancer-gene pair (*g, e*), as the same procedure applies to all enhancer-gene pairs independently. Given an enhancer *e* and gene *g*, Let *y*_*cg*_ be the mRNA expression in cell type *c, x*_*ce*_ be the enhancer activity (e.g., mC or ATAC). Let *C* be the number of cell types. A linear model associating *g* and *e* can be written as:

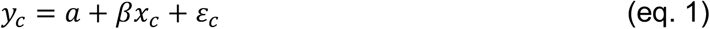

where *c* is the index for cell types, *β* is the association strength, and *ε* is a noise term. In addition, *a* is an intercept term that can be omitted after data centering (*x* and *y* can be pre-centered to ensure *E*[*y*_*c*_] = *E*[*x*_*c*_] = 0). In matrix notation, (eq. 1) can be simply noted as *y* = *βx* + *ε*.

In ordinary least squares (OLS), we assume *ε* is uncorrelated across cell types: *E*[*ε*_*c*_] = 0, *E*[*ε*_*c*_*ε*_*c′*_] = σ^2^*δ*_*c,c′*_. The correlation coefficient *r* = *E*[*xy*]/σ_*x*_σ_*y*_ is then a measure of the linear association, and it has an associated p-value calculated using the t distribution. Alternatively, inference can be performed by permutation analysis to get an empirical p-value.

However, in our case we have correlated noise: *E*[*ε*_*c*_*ε*_*c′*_] = *Ω*_*cic′*_, which reflects the correlation between cell types due to gene co-expression. That is, *Ω*_*c,c′*_ represents the background of correlated variability in gene expression due to the hierarchical structure of cell types in complex tissues. We can estimate the correlation using the genome-wide covariance, 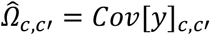. In this case, generalized least squares^16^ (GLS) can be used to give an estimate of the coefficient *β*. This corresponds to transforming the variables *x, y* from the original basis (cell types/metacells, denoted *c*) to an decorrelated basis (denoted *r*), and then performing OLS on the decorrelated variables.

We first use singular value decomposition (SVD) to decompose the mean-subtracted gene expression matrix, *y*_*cg*_ = ∑_1_ *⋃*_*cr*_*S*_*rr*_*V*^*T*^_*rg*_, where *r* = *min*(*c, g*). Defining *Z* = *US*, we have *Ω* = *ZZ*^*T*^. Multiplying both sides of (eq. 1) by *Z*^−1^ = *S*^−1^ *U*^*T*^ corresponds to a transformation from correlated to decorrelated (or whitened) basis:

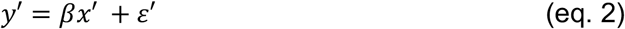

where *y′* = *Z*^−1^*y, x′* = *Z*^−1^*x*, and *ε′* = *Z*^−1^*ε*. The noise term is now uncorrelated, because

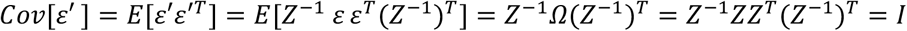

where *I* is the identity matrix. We can therefore use the correlation coefficient and its associated test statistics on transformed data *y′*and *x′*, as in the case of OLS.

### Expected range of correlation coefficients for independent variables

Here we provide theoretical justification on why we expect the range of correlation coefficients 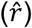 to scale as 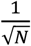, as seen in Fig. 2j and Supplementary Fig. 7b, where *N* is the number of metacells.

Let *X* and *Y* be two independent random variables. Let *x*_*i*_ and *y*_*i*_ be independent and identically distributed samples of *X* and *Y*, where *i* ∈ {1,2, …, *N*}. In our case, *N* represents the number of metacells, and *x*_*i*_ and *y*_*i*_ are the transcriptomic and epigenetic signals for a given enhancer-gene pair for metacell *i*. We require *X* and *Y* to be independent of each other as they are unlinked, and *x*_*i*_ and *y*_*i*_ be independent samples as different metacells are also independent observations of *X* and *Y*, such as in the case of null distribution created by shuffling cells.

To simplify the notation, we assume *E*[*X*] = *E*[*Y*] = 0, as the mean does not affect correlation coefficient *r*. We also assume *X* and *Y* are symmetric, as in the case of normal distribution. It is obvious that *r*(*X,Y*) = 0. However, we are interested in how the variance of 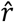 depends on *N*, where 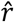 is the sample estimate of *r* by {*x*_*i*_} and {y_*i*_}.

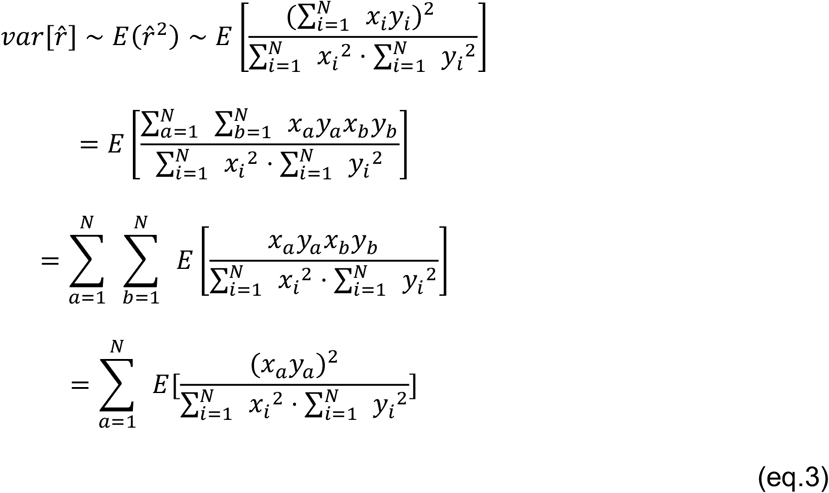

The last equality holds, as only non-interaction terms (*a* = *b*) are nonzero. Moreover, as (*x*_*a*_*y*_*a*_)^2^ 603 equivalent for different *a* = {1… *N*}, the above summation can be further simplified as:

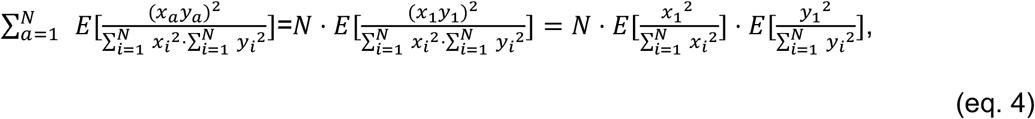

where 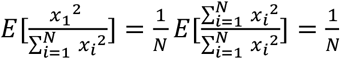, due to the symmetry among indices. Therefore, we finally arrive at

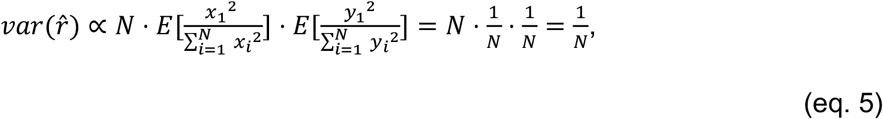

and thus the range of the distribution goes as 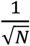.

## Supplementary tables

**Table S1.** A list of putative enhancers (cCREs; n=233,524 in total)

**Table S2.** Significant linked enhancer-gene pairs by mCG-RNA correlation

**Table S3.** Significant linked enhancer-gene pairs by ATAC-RNA correlation

**Table S4.** Cell type correspondence between this study and Ref^6^

## Acknowledgements

We gratefully acknowledge members of the Mukamel, Ecker, Ren, and Zeng laboratories collaborators within the BRAIN Initiative Cell Census Network (BICCN). This work was funded by the NIH BRAIN Initiative (RF1 MH120015 to E.A.M.; U19MH114830 to H.Z.; U19MH121282 to J.R.E.; J.R.E is an Investigator of the Howard Hughes Medical Institute) and by CZI Collaborative Computational Tools for the Human Cell Atlas (to E.A.M.).

## Author contributions

EAM and FX designed the study. ZY, BT, and HZ generated scRNA-seq data. HL, AB, MMB, JDL, CL, JRN, APD, ACR and JRE generated DNA methylation (snmC-Seq) data. HL, MMB, YEL, JDL, APD, OP, SP, and BR generated snATAC-Seq data. FX led the computational analysis. FX and EA developed code and performed analysis. FX, EA, and EAM wrote and edited the manuscript. All authors approved the manuscript.

## Data availability

The scRNA-seq, snmC-seq, and snATAC-seq datasets from the mouse primary motor cortex are generated by BICCN (RRID:SCR_015820) as reported previously^6^. The data can be accessed via the NeMO archive (RRID:SCR_002001) at accession: https://assets.nemoarchive.org/dat-ch1nqb7. Genome browser: https://brainome.ucsd.edu/BICCN_MOp. The chromatin contact data generated by snm3C-seq is downloaded from publicly available files (Ref^17^; https://salkinstitute.app.box.com/s/fp63a4j36m5k255dhje3zcj5kfuzkyj1).

## Code availability

Analysis scripts used for this paper are at https://github.com/FangmingXie/scf_enhancer_paper.

SingleCellFusion is available at https://github.com/mukamel-lab/SingleCellFusion.

The code for ABC model analysis is downloaded from its github repository (https://github.com/broadinstitute/ABC-Enhancer-Gene-Prediction; Ref^5^).

The code for CICERO analysis is downloaded as an R package (https://www.bioconductor.org/packages/release/bioc/html/cicero.html; Ref^8^).

## References

1. Liu, H. et al. DNA methylation atlas of the mouse brain at single-cell resolution. Nature 598, 120–128 (2021).

2. Li, Y. E. et al. An atlas of gene regulatory elements in adult mouse cerebrum. Nature 598, 129–136 (2021).

3. Gasperini, M., Tome, J. M. & Shendure, J. Towards a comprehensive catalogue of validated and target-linked human enhancers. Nat. Rev. Genet. 21, 292–310 (2020).

4. Gasperini, M. et al. A Genome-wide Framework for Mapping Gene Regulation via Cellular Genetic Screens. Cell 176, 1516 (2019).

5. Fulco, C. P. et al. Activity-by-contact model of enhancer–promoter regulation from thousands of CRISPR perturbations. Nat. Genet. 51, 1664–1669 (2019).

6. Yao, Z. et al. A transcriptomic and epigenomic cell atlas of the mouse primary motor cortex. Nature 598, 103–110 (2021).

7. Yao, Z. et al. A taxonomy of transcriptomic cell types across the isocortex and hippocampal formation. Cell 184, 3222–3241.e26 (2021).

8. Pliner, H. A. et al. Cicero Predicts cis-Regulatory DNA Interactions from Single-Cell Chromatin Accessibility Data. Mol. Cell 71, 858–871.e8 (2018).

9. Cusanovich, D. A. et al. A Single-Cell Atlas of In Vivo Mammalian Chromatin Accessibility. Cell 174, 1309–1324.e18 (2018).

10. Zhu, C. et al. An ultra high-throughput method for single-cell joint analysis of open chromatin and transcriptome. Nat. Struct. Mol. Biol. 26, 1063–1070 (2019).

11. Corces, M. R. et al. The chromatin accessibility landscape of primary human cancers. Science 362, eaav1898 (2018).

12. Trevino, A. E. et al. Chromatin accessibility dynamics in a model of human forebrain development. Science 367, eaay1645 (2020).

13. Ma, S. et al. Chromatin Potential Identified by Shared Single-Cell Profiling of RNA and Chromatin. Cell 183, 1103–1116.e20 (2020).

14. Gorkin, D. U. et al. An atlas of dynamic chromatin landscapes in mouse fetal development. Nature 583, 744–751 (2020).

15. Sarropoulos, I. et al. Developmental and evolutionary dynamics of cis-regulatory elements in mouse cerebellar cells. Science 373, (2021).

16. Aitken, A. C. On Least Squares and Linear Combination of Observations. Proceedings of the Royal Society of Edinburgh 55, 42–48 (1936).

17. Lee, D.-S. et al. Simultaneous profiling of 3D genome structure and DNA methylation in single human cells. Nat. Methods 16, 999–1006 (2019).

18. Serwach, K. & Gruszczynska-Biegala, J. STIM Proteins and Glutamate Receptors in Neurons: Role in Neuronal Physiology and Neurodegenerative Diseases. Int. J. Mol. Sci. 20, 2289 (2019).

19. Schoenfelder, S. & Fraser, P. Long-range enhancer-promoter contacts in gene expression control. Nat. Rev. Genet. 20, 437–455 (2019).

20. Endersby, J. Lumpers and splitters: Darwin, Hooker, and the search for order. Science 326, 1496–1499 (2009).

21. Welch, J. D. et al. Single-Cell Multi-omic Integration Compares and Contrasts Features of Brain Cell Identity. Cell 177, 1873–1887.e17 (2019).

22. Luo, C. et al. Single nucleus multi-omics links human cortical cell regulatory genome diversity to disease risk variants. bioRxiv 2019.12.11.873398 (2019) doi:10.1101/2019.12.11.873398.

23. Baran, Y. et al. MetaCell: analysis of single-cell RNA-seq data using K-nn graph partitions. Genome Biol. 20, 206 (2019).

24. Nettleton, D., Hwang, J. T. G., Caldo, R. A. & Wise, R. P. Estimating the number of true null hypotheses from a histogram of p values. J. Agric. Biol. Environ. Stat. 11, 337 (2006).

25. Yin, Y. et al. Impact of cytosine methylation on DNA binding specificities of human transcription factors. Science 356, eaaj2239 (2017).

26. Daigle, T. L. et al. A Suite of Transgenic Driver and Reporter Mouse Lines with Enhanced Brain-Cell-Type Targeting and Functionality. Cell 174, 465–480.e22 (2018).

27. Graybuck, L. T. et al. Enhancer viruses for combinatorial cell-subclass-specific labeling. Neuron 109, 1449–1464.e13 (2021).

28. de Boer, C. G. et al. Deciphering eukaryotic gene-regulatory logic with 100 million random promoters. Nat. Biotechnol. 38, 56–65 (2020).

29. Kent, W. J. et al. The human genome browser at UCSC. Genome Res. 12, 996–1006 (2002).

30. He, Y. et al. Improved regulatory element prediction based on tissue-specific local epigenomic signatures. Proc. Natl. Acad. Sci. U. S. A. 114, E1633–E1640 (2017).

31. Quinlan, A. R. & Hall, I. M. BEDTools: a flexible suite of utilities for comparing genomic features. Bioinformatics 26, 841–842 (2010).

32. Amemiya, H. M., Kundaje, A. & Boyle, A. P. The ENCODE Blacklist: Identification of Problematic Regions of the Genome. Sci. Rep. 9, 9354 (2019).

33. Zeng, H. & Sanes, J. R. Neuronal cell-type classification: challenges, opportunities and the path forward. Nat. Rev. Neurosci. 18, 530–546 (2017).

34. Traag, V. A., Waltman, L. & van Eck, N. J. From Louvain to Leiden: guaranteeing well-connected communities. Sci. Rep. 9, 5233 (2019).

35. Benjamini, Y. & Hochberg, Y. Controlling the false discovery rate: a practical and powerful approach to multiple testing. J. R. Stat. Soc. Series B Stat. Methodol. 289–300 (1995).

36. Imakaev, M. et al. Iterative correction of Hi-C data reveals hallmarks of chromosome organization. Nat. Methods 9, 999–1003 (2012).

37. Abdennur, N. & Mirny, L. A. Cooler: scalable storage for Hi-C data and other genomically labeled arrays. Bioinformatics 36, 311–316 (2020).

